# Transcriptome profiling and comparison of *Rhinanthus major* and *Rhinanthus minor* reciprocal F_1_ hybrids during seed stratification and germination

**DOI:** 10.1101/2021.04.06.438703

**Authors:** Khaled Mirzaei, Renate A. Wesselingh

## Abstract

- Background and Aims Germination is a vital stage in a plant’s life cycle, and a different germination behavior of offspring in comparison to their parents can have fitness consequences. In studies on hybridization between *Rhinanthus minor* and *R. major*, low germination rates of F_1_ hybrids with *R. major* as the maternal parent have often been reported. In contrast, the F_1m_ hybrid, with *R. minor* as the maternal parent, germinates readily and rapidly.
- Methods In order to find the cause of this difference, we used RNA-Seq to obtain transcriptome profiles of F_1a_ and F_1m_ seeds during stratification at 4°C and just after germination, after 40 days of stratification for the F_1m_ seeds and 60 days for the F_1a_ seeds.
- Key Results A comparison of the transcriptome of F_1a_ seeds that had just germinated (60 days) with non-germinated F_1a_ seeds after 40 and 60 days revealed 2918 and 1349 differentially expressed (DE) genes, respectively. For F_1m_ seeds, 958 genes showed differential expression in germinated and non-germinated seeds after 40 days. The DE genes of F_1a_ and F_1m_ hybrids clustered into two separate groups, even though they had the same parents, and no differentially expression was found for plastid genes. Non-germinated F_1a_ seeds had an abundance of enzymes and proteins associated with peroxidase activity, peroxiredoxin activity and nutrient reservoir activity. Expression of genes related to seed germination and seed development increased in non-germinated F_1a_ hybrid seeds between 40 and 60 days of cold stratification. F_1a_ seeds that had germinated showed an upregulation of genes related to the gibberellic acid-mediated signaling pathway and response to gibberellin, along with a low expression of DELLA superfamily.
- Conclusions Although the results demonstrated strong differences in gene expression during stratification between the reciprocal hybrids, we could not identify its cause, since no plastid genes were differentially expressed. It is possible that differences in embryo development after seed formation and before stratification play a role, including epigenetic imprinting.

## Introduction

Natural hybridization is a source of novel genetic material for evolution and it can occur among species due to a lack of reproductive barriers. It is more common among plants than animals (Rieseberg 1997), and although it is rare in some families, in others, such as *Dennstaedtiaceae*, it is very common phenomena (Whitney *et al.* 2010). When hybridization occurs, the fate of the F_1_ hybrid can vary from showing hybrid vigor—which has greatly benefited agriculture—to hybrid breakdown, which can be a key factor in speciation (Rieseberg 1997; Arnold *et al.* 1999; Johansen-Morris and Latta 2006). Hybrid survival can play an important role in determining the effect of hybridization on the future composition of population, given that a hybrid can be a bridge for introgression or become a new species (Stukenbrock 2016; Grant and Grant 2019). By providing opportunities for gene flow between sister species, hybridization can cause a decrease or an increase (reinforcement) of reproductive barriers (Pickup *et al.* 2019).

Most flowering plant species are hermaphrodite, and therefore the formation of reciprocal hybrids is often possible, but these reciprocal hybrids do not always have the same performance (Christopher *et al.* 2019). Asymmetry in the fitness of reciprocal F_1_ hybrids has been reported in many plant species (Tiffin *et al.* 2001; Turelli and Moyle 2007). This type of isolation, known as isolation asymmetry, is different from Dobzhansky–Muller incompatibilities (DMIs) because the reciprocal F_1_ hybrids carry the same autosomal genotype (Turelli and Moyle 2007). The combination of two divergent genomes through hybridization will introduce immediate and profound genetic modifications and remodeling of parental gene expression in the F_1_, which can cause genome shock and have wide effects on species establishment and diversification (Paun *et al.* 2007; Kerbs *et al.* 2017). Due to extrinsic or intrinsic factors, reciprocal hybrids may show differences in fitness and can be different from their parents in the same environment (Paun *et al.* 2007). According to Darwin’s corollary to Haldane’s rule, isolation asymmetry can be an outcome of different relative rates of evolution in the parents of the F_1_ hybrid (Turelli and Moyle 2007).

*Rhinanthus major* Ehrh. and *R. minor* L. (both 2n = 2x = 22: Hambler (1954)) are annual plants in the Orobanchaceae family, occurring in a diverse range of open habitats (Westbury 2004b; Ducarme *et al.* 2010). Both species are self-compatible, but a higher outcrossing rate has been documented for *R. major* (76%) in comparison with *R. minor* (13%; Ducarme and Wesselingh 2013). Both species are hemiparasites that take organic carbon and mineral nutrients from the root systems of their hosts, even though they are able to photosynthesise (Rümer *et al.* 2007). Their roots have found to be attached to nearly 50 plants species from 18 families, but the most suitable hosts are members of the Poaceae and Fabaceae (Gibson and Watkinson 1991).

Seed germination as a first stage of plant transition to development phase is composed of a series of steps, from water absorbance to embryo development and digestion of the starch and protein reserves of the seed (Bentsink and Koornneef 2008). If seed germination is mainly controlled by dormancy (primary or secondary dormancy), this can provide the best timing for plant growth and development to reduce extrinsic mortality (Hoyle *et al.* 2015).

Differences in seed dormancy within the same species can place the earliest and the latest plant in different environments with different challenges and this can provide an opportunity for adaptive divergence (Donohue *et al.* 2005; Hoyle *et al.* 2015). Naturally, seeds of *Rhinanthus* species have physiological dormancy that can be broken by several weeks of cold stratification (Westbury 2004; Ter Borg 2005; Marin *et al.* 2019). Under laboratory conditions, a significant difference in germination rate has repeatedly been observed between the reciprocal F_1_ hybrids. The germination rate of hybrids formed on *R. major* (F_1a_ hybrids) is only 5-30%, whereas hybrids formed with *R. minor* as the maternal species (F_1m_ hybrids) is nearly 100%, often even better than the parental species (Kwak 1979; Campion-Bourget 1980; Natalis and Wesselingh 2012; Ducarme and Wesselingh 2013).

However, this difference was not observed in a field experiment (Wesselingh *et al.* 2019), which seems to indicate that the experimental conditions used for germination in the laboratory are not representative for the germination behaviour of the hybrids under more natural conditions. Marin *et al.* (2019) have shown in *R. minor* that seed features related to quality such as germination capacity and seed vigor can play important role in plant emergence, establishment and performance. They also found that elongation and growth of *R. minor* embryo continued after seed dispersal. Anatomical studies of *R. major* showed that its seed contained a well-developed embryo before the start of stratification (Tiagi 1966).

Gene expression undergoes important changes during seed development, and comparative studies of transcriptomes of seeds during dormancy and germination have helped scientists to understand the processes that are taking place. Among expression profiling technologies, RNA sequencing (RNA-Seq), a member of next–generation sequencing (NGS) technologies, is a comprehensively informative technique to monitor wide transcriptional changes during various stages of plant life. Therefore, we used RNA-Seq to study the genetic basis and transcriptional changes during germination of F_1a_ and F_1m_ hybrids.

## Material and methods

### Plant materials and RNA extraction

Hybrid seeds were produced by reciprocal hand pollination on plants grown from seed in the greenhouse in 2019, applying *R. major* pollen onto *R. minor* stigmas after emasculation to produce F_1m_ hybrid seed, and *R. minor* pollen onto *R. major* stigmas for F_1a_ hybrid seeds. Seeds were collected from dry capsules in June-July 2019 and stored at room temperature. Thirty seeds in three replicates from each cross type were placed in petri dishes for germination in a refrigerator (± 4°C) in October 2019. We checked the dishes regularly for germination, which was scored as soon as the radicle was visible. We sampled germinated seeds at an early stage, with less than 1 mm of the radicle protruding from the seed, and sampled non-germinated seeds at the same time, after 40 days and 60 days of stratification. RNA was extracted from seeds using the QIAGEN RNeasy Mini Kit (3 seeds per sample): non-germinated F_1m_ (F1m_40_NG) and F_1a_ seeds (F1a_40_NG) 40 days after the start of stratification, germinated F_1m_ seeds after 40 days (F1m_40_G), germinated F_1a_ seeds after 60 days (F1a_60_G), and non-germinated F_1a_ seeds after 60 days (F1a_60_NG). The columns used for RNA extraction were washed again by TE buffer and centrifuged for 5 minutes at 10000 rpm to remove the DNA. The retrieved DNA was used for PCR (20 μl final volume, 2 μl of 10 X buffer, 0.4 mM dNTPs, 1.5 mM MgCl2, 2.0 U Taq polymerase, 5 pM forward primer and 5 pM reverse primer) with species-specific primers (5’CACCCTGATTTCTCTTTCTTCAA, 5’TTAAGACCCCATAAAAAGGAGGA) and obtained PCR products were digested by RsaI enzyme to confirm if they were indeed hybrid (Wesselingh *et al.* 2019; Mirzaei and Wesselingh 2021). Finally, the RNA of 15 hybrid samples was sent to GENEWIZ for sequencing on Illumina NovaSeq (2*150 bp) platform.

### De novo assembly and sequence annotation

Preliminary quality checks of all sequenced samples were performed using FastQC and quality trimming and adaptor removal were done using Trimmomatic-0.39 (Bolger *et al.* 2014). *In silico* normalization of reads was done using the normalization option in Trinity package and *de novo* transcriptome assembly was performed using three assemblers: Trinity (Grabherr *et al.* 2011), Trans-ABySS (Robertson *et al.* 2010) and SPAdes-rna (Bushmanova *et al.* 2019). A first appraisal of the quality of the assembled transcriptomes was performed by estimating representation read counts in each assembly using bowtie2 (Langmead and Salzberg 2012). Transcriptome completeness was explored using BUSCO v.4.1.2 (Benchmarking Universal Single Copy Orthologs) (Simão *et al.* 2015) to obtain the percentage of single-copy orthologues represented. Plotting of the BUSCO results was performed using the ggplot package in R v4.03 (R Core Team 2020). Functional annotation of transcriptome was conducted using the Trinotate (Bryant *et al.* 2017). Prediction of coding regions in transcripts and open reading frames (ORFs) were performed by TransDecoder v5.5.0 (Grabherr *et al.* 2011). UniProtKB/Swiss-Prot and plant NCBI NR (NCBI non-redundant protein database) databases were used for homology searches (Ye *et al.* 2006) and HMMER v.3 (Finn *et al.* 2011) and Pfam (Punta *et al.* 2012) were used for protein domain identification. Signal peptide predictions was obtained using signalP v.445 (Petersen *et al.* 2011) and transmembrane regions were predicted using the tmHMM v.246 server (Krogh *et al.* 2001). Ribosomal RNA genes were detected with RNAMMER v.1.247 (Lagesen *et al.* 2007) and finally annotation outputs were loaded into a Trinotate SQLite Database.

### Transcript abundance and differential expression analysis

To estimate transcript abundance, we used the alignment-based RSEM as abundance estimation method. Therefore, we used bowtie2 for alignment, and the sorted alignment file in BAM format was generated by SAMtools-1.3.1 (Li *et al.* 2009) and used for RSEM program (Li and Dewey 2011). Finally, a matrix of normalized expression values were estimated with the abundance_estimates_to_matrix.pl script in the Trinity package. The resulting matrix of normalized expression values was fed into the count_matrix_features_given_MIN_TPM_threshold.pl script for ExN50 analysis. The ExN50 statistics was calculated by Trinity accessory scripts contig_ExN50_statistic.pl and plotted with ggplot in R. Differential expression was preformed using adapted pipeline in Trinity which included Bioconductor v3.4, edgeR v4.0 (Robinson *et al.* 2010), Limma (Ritchie *et al.* 2015), ctc (Lucas and Jasson 2006), Biobase (Huber *et al.* 2015), gplots (Warnes *et al.* 2009). The transcript differential expression analysis was performed on the matrix of raw read counts using the edgeR R package. The false discovery rate raw *p*-values were adjusted for multiple comparisons by the Benjamini-Hochberg method (Haynes 2013). A false discovery rate [FDR] < 0.05 and |log2FC| ≥ 2 (positive or negative used for over- or under-expression, respectively) were used as criteria for identifying significant differences in expression. Gene set enrichment analyses of GO terms were conducted on each set of differentially expressed transcripts to determine over-represented functional pathways. GO enrichment analysis of differentially expressed (DE) genes was performed using Fisher exact test, P-value ≤0.05 and the hypergeometric Fisher exact test (P < 0.05) and Benjamini (FDR < 0.05) were used to detect statistically significant enrichment of the KEGG pathway (Kanehisa *et al.* 2016).

## Results

### Germination and RNA-sequencing

In the F_1m_ hybrid seeds, germination started after 34 days and after 70 days 90% of seeds had germinated. On the contrary, F_1a_ hybrid seeds started to germinate after 39 days and even after 120 days in the refrigerator the germination rate was still under 17%. The mRNA libraries of the five treatments generated over ~296 million raw paired-end reads, out of which ~263 million high quality paired-end reads (Q≥ 20) remained with ~17.53 million reads per sample. The final lengths of remaining reads were between 36 and 151bp, with a GC content of approximately 48% in all samples (Supplementary DataTable S1).

### Assembly

The Trinity assembly, with more complete reads (91.8%) and a higher representation of sequenced reads (97.53%) had a better quality in comparison with the two other assemblies (Table 1). The total length of assembled reads, number of transcripts and the N50 index of Trinity assembly were also higher than in the Trans-ABySS and SPAdes-rna assemblies. Therefore we used the Trinity assembly for further analysis.

**Table 1.** Summary of assembly statistics generated by various pipelines. BUSCO Eukarya database OrthoDB v.10 busco genes.

| Workflow | Trinity | Trans-ABYSS | SPAdes-rna |
| --- | --- | --- | --- |
| Representation of RNA-seq reads | 97.53% | 82.8% | 74.59% |
| **BUSCO Results** |  |  |  |
| Complete | 91.8% | 87.3% | 58.1% |
| Fragmented | 3.1% | 8.6% | 22.3% |
| Missed | 5.1% | 4.1% | 19.6 |
| **General assembly metrics** |  |  |  |
| Length, Mbp | 306 | 298 | 163 |
| Number of transcripts | 356379 | 317485 | 169473 |
| N50, bp | 1558 | 1461 | 1272 |

The Trinity assembly contained a total of 356379 transcripts (206,255 trinity genes) with an average length of 861bp and N50 value of 1,558bp (Table 1). The lengths of the assembled transcripts ranged from 200 to 22,700 bp and about 74.34% of the transcripts were in the range of 201–1000 bp (~1 kb), 23.98% transcripts were 1001–4000 bp (1.1 to 4.0 kb) and 1.68% was longer than 4001 bp (>3 kb) (Supplementary Data Fig. S1). Plotting transcript expression (Ex) against ExN50 value-as another overall quality checking of assembly-revealed that saturation point of the assembly was at 92% of the total expression with length of 1884 bp (Supplementary Data Fig. S2). According to ExN50 value, deeper sequencing is very unlikely to provide longer reads for our transcriptomes (Supplementary Data Fig. S2).

### Annotation

In total 173,638 ORFs were predicted, including: 92241 (53.12%) as complete ORFs contained a starting codon for methionine and ending stop codon, 32193 (18.54%) as 5’ partial ORF which lacked the start codon, 18825 (10.84%) as 3’-partial ORF and 30329 (17.46%) as internal ORF which were partial at both 5’ and 3’. Nearly 78% of the ORFs and 66% of the transcripts were matched with the Swiss-Prot database and 58% with the plant NR database. Homologous genes with high probability scores were found for 58.03% (100775) of predicted ORFs. Taxonomic homology search of ORFs revealed that the highest similarity (5%) of the ORFs matched with *Capsicum annuum* (Solanaceae) and *Gossypium raimondii* (Malvaceae) and only 2% (2157) with *Striga asiatica* from Orobanchaceae family (Supplementary Data Table S2).

### Differential gene expression analysis

A preliminary comparison of replicates across all samples was performed using a PCA on normalized read counts (Fig. 1). The first principal component, which explained 27.80% of the observed variation, clearly separated the two reciprocal hybrids. The second component (13.5%) allowed to distinguish between germinated and non-germinated seeds of the F_1a_ hybrid. A clear separation between 40 and 60 days of stratification for the non-germinated F_1a_ hybrid seeds is visible on the third principal axis (11.93%). Correlation analysis of differentially expressed (DE) genes clearly revealed two groups, again corresponding to the two reciprocal hybrids (Fig. 2).

**Fig. 1.**
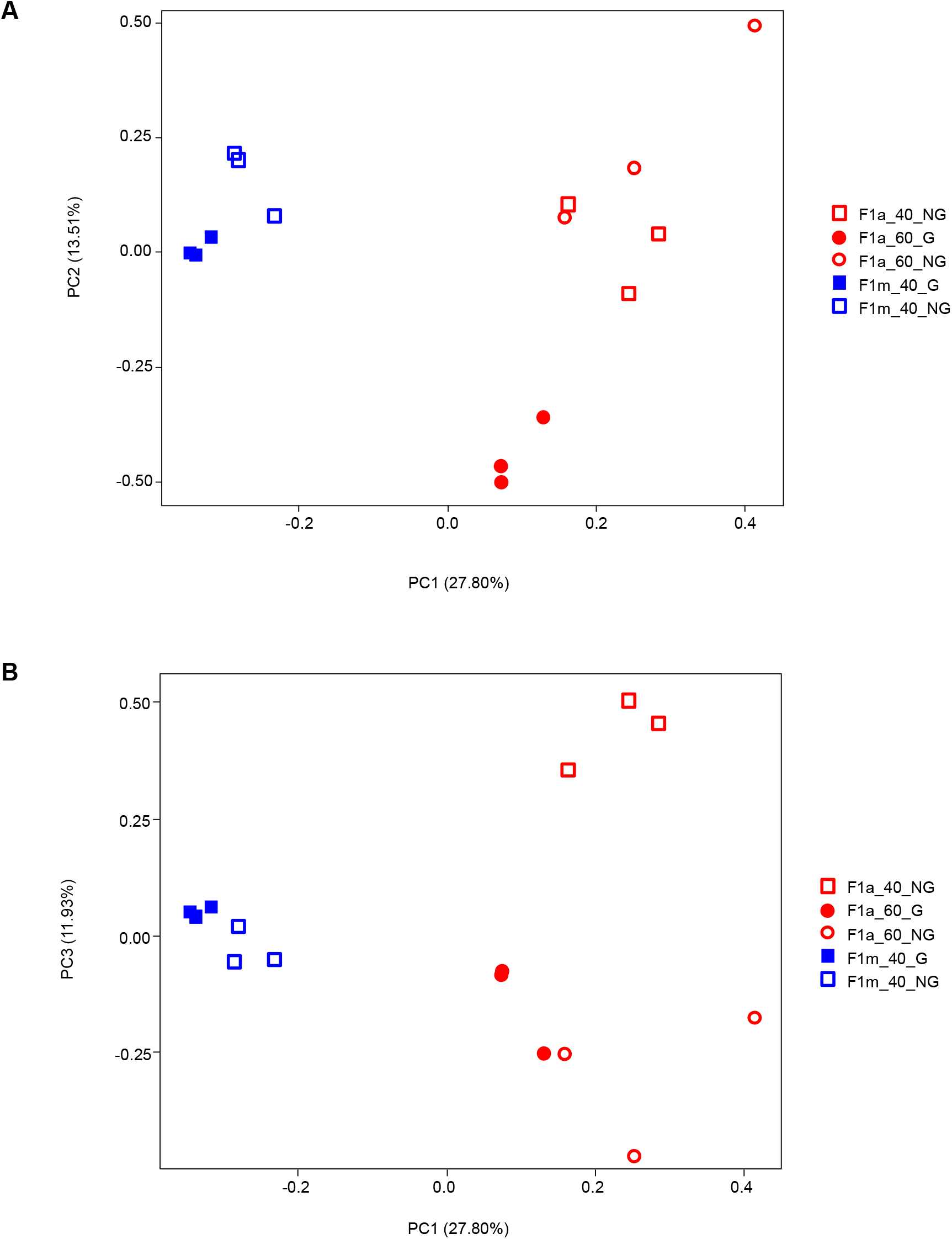
Principal component analysis of RNA-seq samples of *Rhinanthus* hybrid seeds using variance-stabilized estimated raw counts. a) PC1-PC2, b) PC1-PC3

**Fig. 2.**
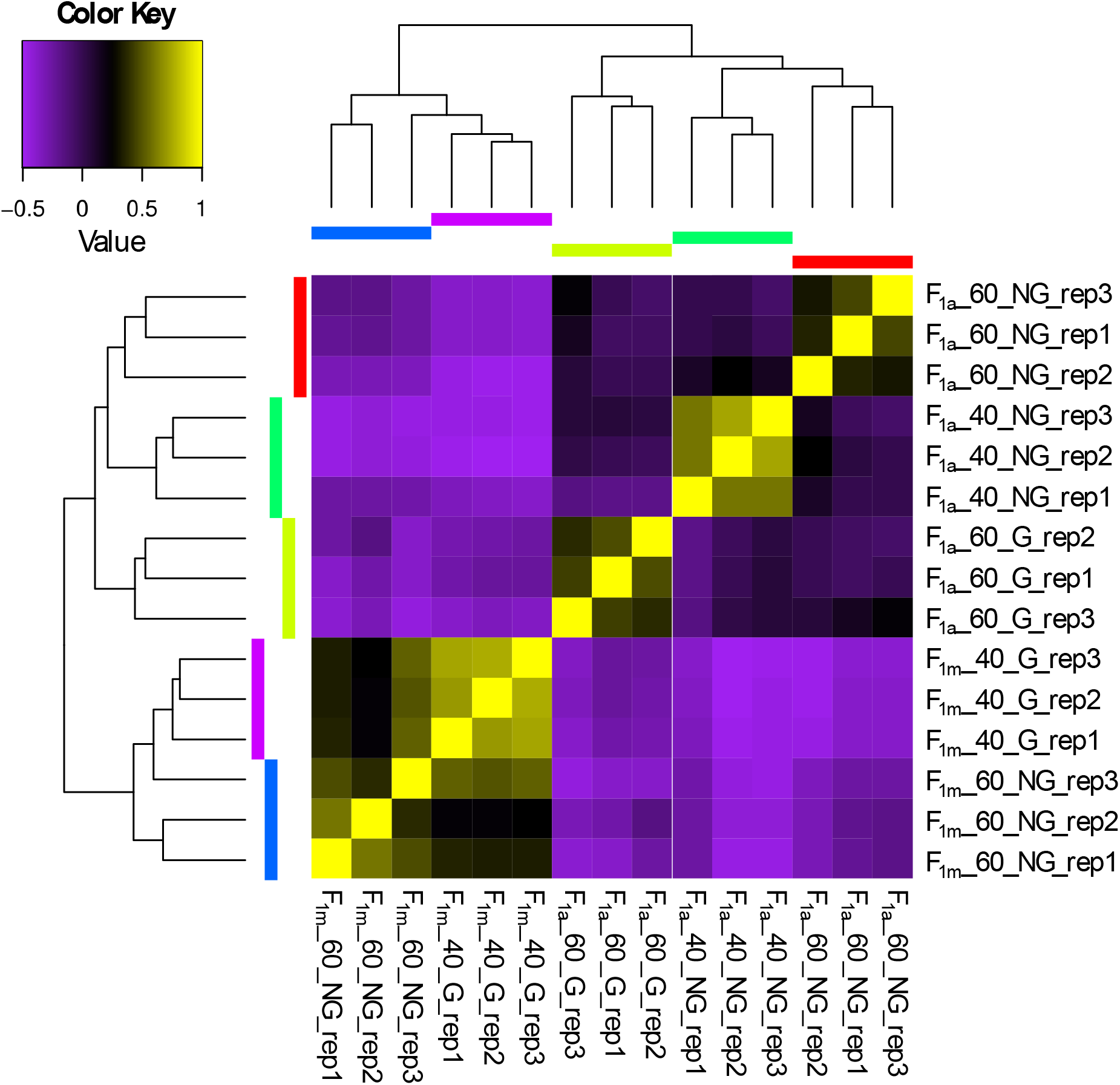
Comparison among replicates in each group of germinated and non-germinated seeds of reciprocal F_1_ hybrids between *Rhinanthus major* (a) and *R. minor* (m).

The highest number of DE genes (4165) was detected between non-germinated F_1a_ seeds at 40 days and germinated F_1a_ seed at 60 days with respectively 1634 and 2531 upregulated genes (Table 2; Fig. 3; Supplementary Data Tables S5). Forty-three upregulated genes such as phytochrome B (phyB), NAC domain-containing protein, and oil body-associated proteins were shared among non-germinated seeds, while 399 shared upregulated genes were observed in germinated seeds (Fig. 3).

**Table 2.** Number of differentially expressed transcripts in three treatment comparisons ([FDR] < 0.05 and |log2FC| ≥ 2) in F_1_ hybrids between *Rhinanthus major* (a) and *R. minor* (m), the letter indicating the maternal parent. NG = non-germinated seeds, G = germinated seeds, after 40 or 60 days on wet filter paper at 4°C.

| Treatment | F<sub>1a_60</sub>_NG vs F<sub>1a_60</sub>_G |  | F<sub>1a_40</sub>_NG vs F<sub>1a_60</sub>_G |  | F<sub>1m_40</sub>_NG vs F<sub>1m_40</sub>_G |  | F<sub>1a_60</sub>_NG vs F<sub>1a_40</sub>_NG |  |
| --- | --- | --- | --- | --- | --- | --- | --- | --- |
| Number of upregulated genes | 735 | 1158 | 1634 | 2531 | 550 | 868 | 1053 | 1363 |

**Fig. 3.**
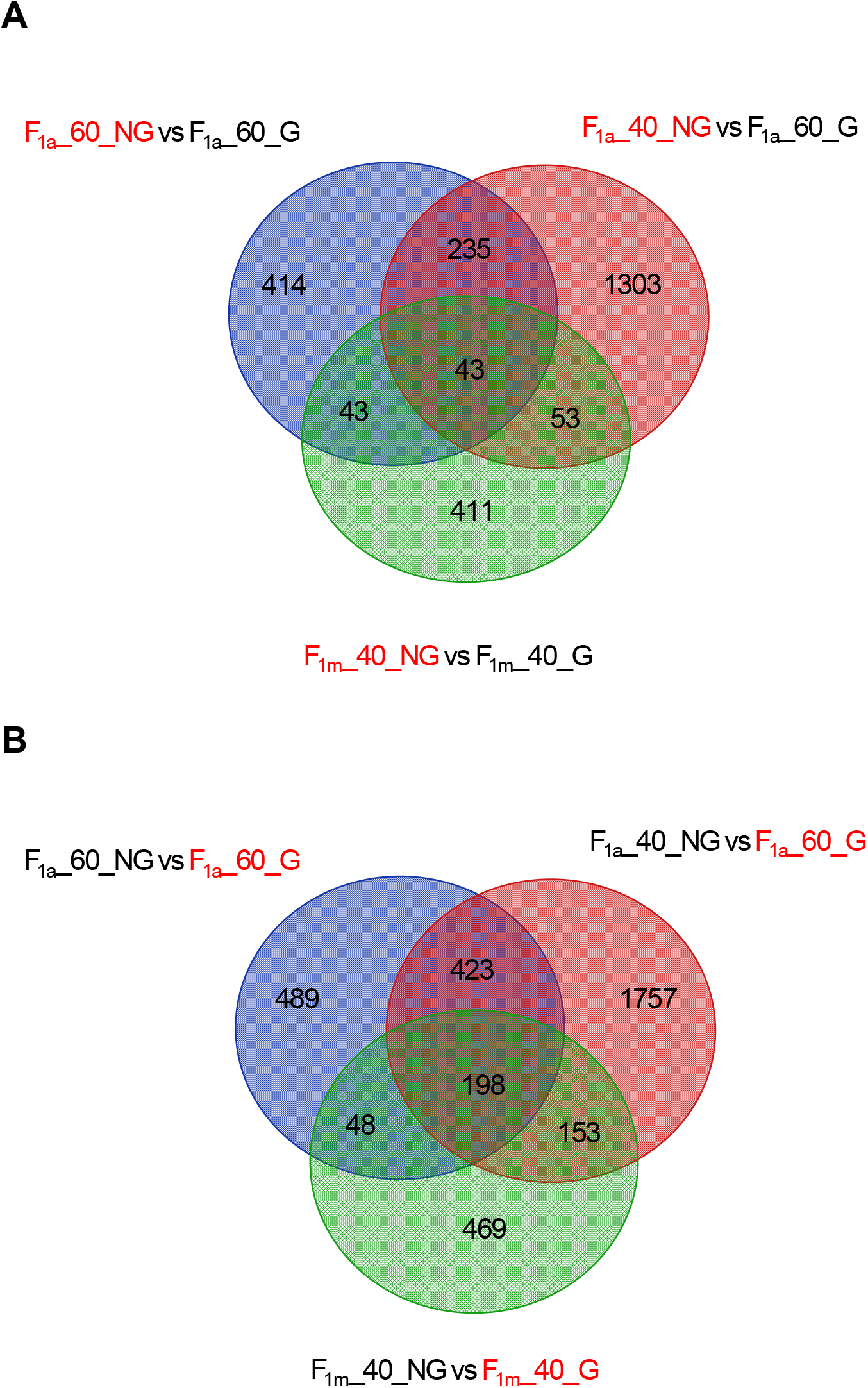
Venn diagram showing the numbers of unique and shared upregulated genes in each comparative analysis between non-germinated (NG) seeds and germinated seeds (G) for a) the non-germinated seeds (F_1a__60_NG, F_1a__40_NG and F_1m__40_NG), and b) the germinated seeds (F_1a__60_G and F_1m__40_G).

In total 176380 (49.50%) and 167885 (47.10%) transcripts were assigned to GO terms and KEGG pathways, respectively. GO enrichment analysis of DE genes in the non-germinated seeds assigned them into 71, 81 and 70 significant GO functional groups, respectively, within three main categories: molecular function, biological process, and cellular components (Supplementary Data Tables S3). The GO molecular function category related to nutrient reservoir activity (GO:0045735) and oxidoreductase activity (GO:0016491), with the highest number of upregulated genes that were significantly enriched in F_1a__40_NG vs F_1a__60_G and in F_1a__60_G vs F_1a__40_NG (Fig. 4A, 4B). Alcohol dehydrogenase (NAD+) activity (GO:0004022) and oxidoreductase activity (GO:0016491) were found to be significantly enriched in F_1m__40_NG vs F_1m__40_G upregulated genes (Fig. 4C). In terms of biological process, response to salt stress (GO:0009651) and oxidation-reduction process (GO:0055114) were enriched in F_1m__40_NG vs F_1m__40_G, with the highest number of upregulated genes, whereas among the upregulated genes in F_1a__40_NG (Fig. 4A, 4B, 4C) we did not have any candidates in terms of biological process and cellular components. In terms of cellular components, extracellular region (GO:0005576) was shared between F_1a__60_G and F_1m__40_G (Supplementary Data Tables S3).

**Fig. 4.**
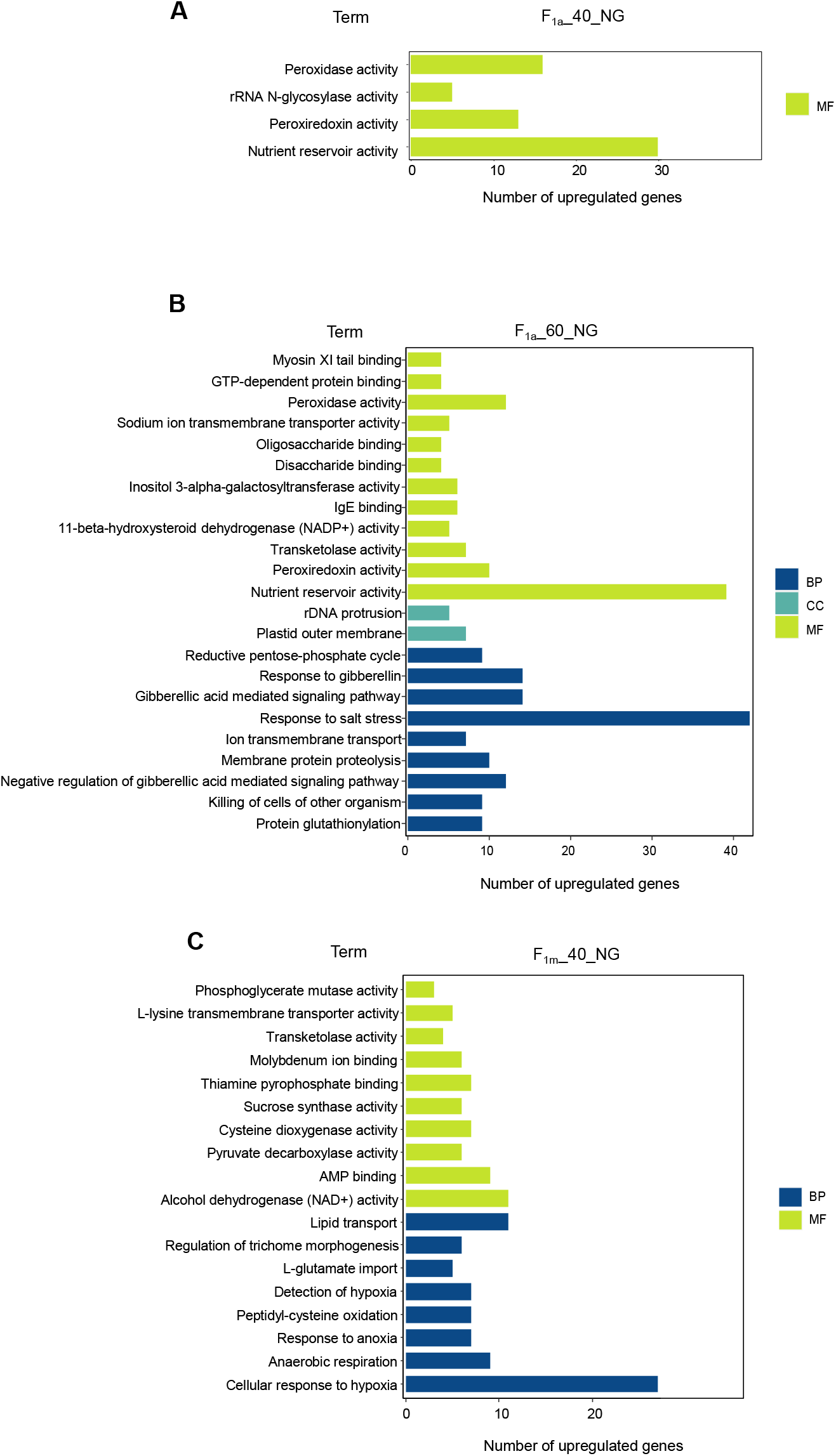
GO pathway enrichment analysis, based on upregulated genes in a) F_1a__40_NG vs F_1a__60_G, b) F_1a__60_NG vs F_1a__60_G and c) F_1m__40_NG vs F_1m__40_G. BP = Biological process, CC= Cell cycle, MF= Molecular function

In total 20 unique KEGG pathways displayed significant changes (P-value ≤0.05) in all compared treatments. “Metabolic pathways”, “plant hormone signal transduction” and “biosynthesis of secondary metabolites” were represented among all the treatments. “Ubiquitin mediated proteolysis” and “amino sugar and nucleotide sugar metabolism” pathways were also common between F_1a__60_NG vs F_1a__60_G and F_1a__40_NG vs F_1a__60_G treatments. “Linoleic acid metabolism” and “starch and sucrose metabolism” pathways were found to be unique in F_1m__40_NG vs F_1m__40_G (Fig. 5; Supplementary Data Table S4).

**Fig. 5.**
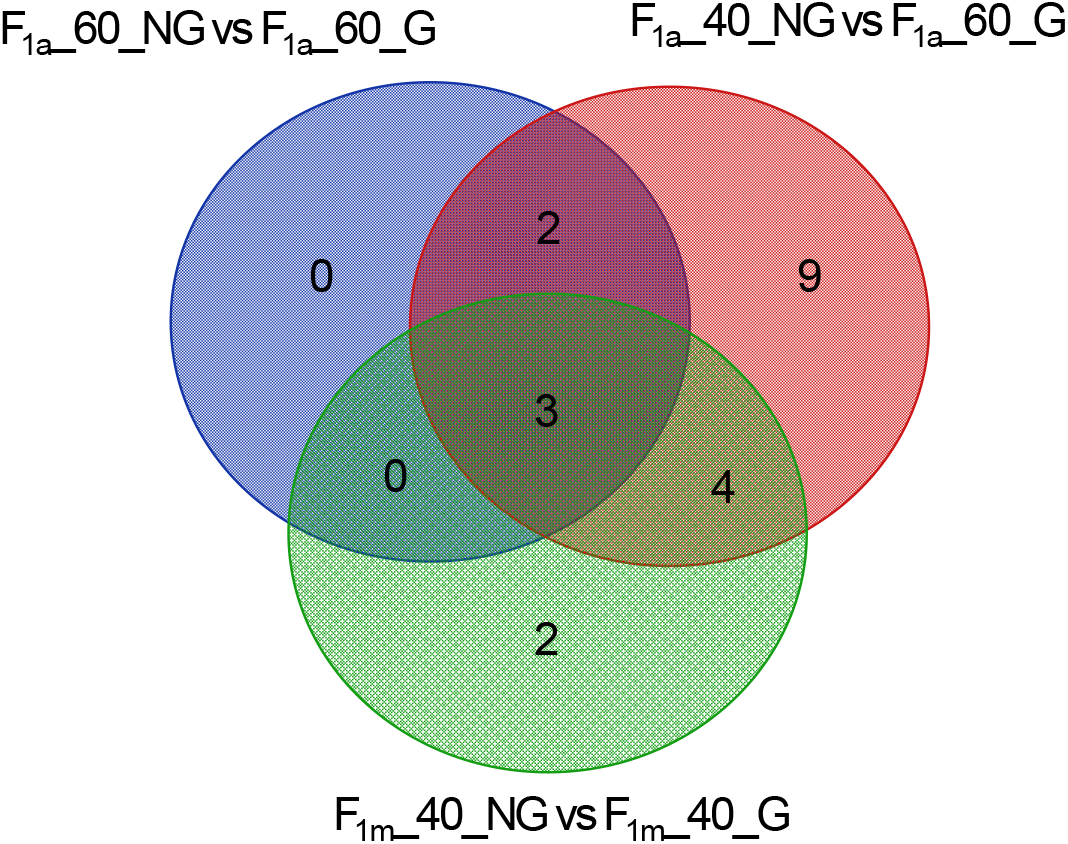
Venn diagram showing the number of unique and shared KEGG pathways related to DE genes in germinated and non-germinated reciprocal F_1_ hybrid seeds.

## Discussion

Overall, the gene expression profiles differed markedly between the two reciprocal hybrids, both before and after germination. Within the F_1a_ hybrids, the differences in gene expression between germinated and non-germinated seeds were stronger than for the F_1m_ hybrid seeds, and they also changed over the course of the stratification period.

### DE genes related to seed development and osmotic stresses

Among the upregulated genes in the non-germinated F_1a_ seeds in comparison with the germinated seeds, peroxidase activity, peroxiredoxin activity and nutrient reservoir activity were common. Plant peroxidases, mostly known for reactive oxygen species (ROS) metabolism, are broadly active in various stages of plant life, from plant development to seed germination (Syros *et al.* 2005). ROS can cause oxidation damage to cell components, but recently it has been proven that ROS can play a key signaling role during germination or dormancy release (Oracz *et al.* 2007; Sarath *et al.* 2007; El-Maarouf-Bouteau and Bailly 2008). One of the sources of ROS in plant cells is peroxisome (glyoxysome) activity (Sandalio and Romero-Puertas 2015). The most significantly enriched upregulated genes in the non-germinated F_1a_ seeds after 60 days were related to glyoxysome, malate synthase activity and the glyoxylate cycle. Peroxiredoxin (Prx) proteins are types of plant antioxidants that protect lipids, enzymes, and DNA against ROS (Rouhier *et al.* 2001). During late seed development and dormancy in mature seeds they are highly expressed, and by sensing harsh conditions they are involved in the maintenance of dormancy while protecting the embryo from damage caused by ROS (Haslekås *et al.* 2003). It has been shown that Prx genes are up– regulated by ABA and osmotic stresses during dormancy and suppressed by gibberellic acid upon germination (Aalen 1999). A recent study in *Arabidopsis thaliana* has shown that *AtPER1*, a seed◻specific peroxiredoxin, is involving in enhancing primary seed dormancy by eliminating ROS to suppress ABA catabolism and GA biosynthesis (Chen *et al.* 2020). The numbers of upregulated genes related to nutrient reservoir activity, which provides the proteins required for the development or growth of seeds, were high in non-germinated F_1a_ seeds, both after 40 and 60 days, whereas only one gene from this category (*VCL21*) was highly expressed in the non-germinated seeds of the reciprocal F_1m_ hybrid at 40 days.

Additionally, GO enrichment resulted in significant representation of rRNA N-glycosylase activity genes in the non-germinated F_1a_ seeds after 40 days. rRNA N-glycosylase activity is a type of toxic activity which depurinates rRNAs and arrests protein synthesis during translation. They have been widely detected in plants and mainly act as antifungal, antibacterial and antiviral agents (Sharma *et al.* 2004; Zhu *et al.* 2018). Glutathionylation activity protects cells against oxidative stress, especially heavy metal stress, and is also involved in many other processes, including cell cycle and cell differentiation, symbiosis and flowering (Rouhier *et al.* 2008; Gao *et al.* 2009; Yadav 2010). Nine genes related to this activity were upregulated after 60 days in non-germinated F_1a_ seeds. It has been shown that this protein is involved in post-transitional modification under oxidative stress conditions and can act as a redox signaling mechanism for helping the cells to sense and signal harmful stress conditions and trigger appropriate responses against stress (Gao *et al.* 2009). Alongside the common GO term enrichment for upregulated genes in the non-germinated F_1a_ seeds in comparison with the germinated F_1a_ seeds, we found the gibberellic acid-mediated signaling pathway, response to gibberellin and negative regulation of gibberellic acid-mediated signaling pathway GO enrichment only after 60 days in the non-germinated F_1a_ seeds.

### DE genes related to phytohormone signal transduction

Maintenance and release of dormancy depend on the intrinsic balance between abscisic acid (ABA) and gibberellic acid (GA). While the maintenance of dormancy depends on high ABA/GA ratios, release of dormancy implies an increased biosynthesis of GA and degradation of ABA, resulting in low ABA/GA ratios (Kermode 2005). The enrichment pattern in non-germinated F_1a_ seeds after 60 days showed an increase in the abundance of enzymes and proteins associated with the gibberellic acid-mediated signaling pathway and response to gibberellin in comparison to 40 days. In contrast, genes related to the negative regulation of the gibberellic acid-mediated signaling pathway and response to salt stress were highly abundant in non-germinated F_1a_ seeds after 60 days. These genes, like membrane-bound NAC transcription factor (*NTL8*), gibberellin 2-oxidases (*GA2oxs*) and ABSCISIC ACID-INSENSITIVE 5 (*ABI5*) are likely to be linked to the maintenance of dormancy in these seeds (Kim *et al.* 2008; Lo *et al.* 2008; Kim and Park 2008). NAC transcription factors (*NTLs*) are related to the stress response and they have negative regulatory effects on seed germination (Kim *et al.* 2007; Kim and Park 2008). *GA2oxs* through 2β-hydroxylation can inactivate GA and increase the ABA/GA ratio (Sakamoto *et al.* 2004). *ABI5* from bZIP transcription factor family has a key role in ABA signaling and inhibiting seed germination (Finkelstein and Lynch 2000).

All of the enriched KEGG pathways for DE genes in non-germinated F_1a_ seeds after 60 days were shared with the same class after 40 days. After 40 days, however, there were 13 more enriched pathways compared to germinated F_1a_ seeds after 60 days, and they were related to lipid metabolism, arginine biosynthesis and glutathione metabolism. Genes clustered in this pathway were mainly upregulated in both germinated and non-germinated seeds after 60 days. Genes related to the “circadian rhythm” pathway were only enriched in upregulated genes in germinated seeds (F_1m_ after 40 days and F_1a_ after 60 days). Although “metabolic pathways” and “biosynthesis of secondary metabolites” were also quite abundant in the other treatments, here we mainly focused on “plant hormone signal transduction” pathway.

Phytohormones like GAs promote germination but ABA and Auxin (Aux) are hormones known to induce and maintain seed dormancy (Liu *et al.* 2013; Tuan *et al.* 2018). In germinated F_1a_ seeds, upregulation of Auxin transporter protein 1 (*AUX1*) and Auxin/Indole-3-Acetic Acid (*Aux/IAA*) was detected, while in non-germinated F_1a_ seeds an upregulation of Transport inhibitor response 1 (*TIR1*) was observed. Studies have shown that Aux/IAA proteins act as positive regulators during seed germination and *TIR1* act as negative regulator for germination (Liu *et al.* 2013; Hussain *et al.* 2020). Aux/IAA protein abundance during germination will promote germination through the inhibition of *ABI3* transcription (Hussain *et al.* 2020). Upregulation of *ABI5* was observed in all of the non-germinated seeds and in non-germinated F_1a_ seeds at 40 days there was an abundance of pyrabactin resistance 1(PYR1)/PYR1-like2 (*PYL2*), *PYL4* and *GAI* with *RGA* (members of the DELLA superfamily). *PYL* receptors are in the ABA signaling pathway and DELLA proteins are key negative regulators of the GA signaling pathway. It has been shown that a high abundance of these genes can induce dormancy and prevent germination (Tyler *et al.* 2004; Tuan *et al.* 2018). The high abundance of *ABI5* and members of the DELLA superfamily in non-germinated seeds could be the main reason of low germination rate of F_1a_ seeds, since we also detected a high expression of *ABI5* in non-germinated F_1m_ seeds after 40 days.

### Maternal effects on F_1_ germination

Large differences in gene expression were observed when we compared F_1_ germinated and non- germinated seeds with different maternal parents in comparison with groups sharing the same maternal parent species. The reciprocal F_1_ seeds differed in the number of days to germination and in the final germination percentage, and this was also conspicuous in the gene expression analysis, which showed clear maternal effects, leading to a prolonged dormancy in the hybrids with *R. major* as the maternal parent.

We only found differential gene expression for nuclear genes, not for mitochondrial or chloroplast genes. Dormancy, germination and seedling establishment can be maternally controlled through maternal tissues surrounding the embryo, such as the endosperm and seed coat (Debeaujon *et al.* 2007; Piskurewicz *et al.* 2016). Maternal effects can be caused by the maternal environment and by the maternal genome (Chiang *et al*. 2011; Fernández Farnocchia *et al*. 2019). The parents of our F_1_ hybrids were grown simultaneously in the same conditions and with the same host plant species, so the observed maternal effects are most likely maternal genetic effects. Even in germinated seeds, we found strong differences in gene expression between the reciprocal hybrids, which indicates the potential long-term nature of these maternal effects.

Although differences in gene expression patterns were clear between the reciprocal hybrids, we cannot yet pinpoint which pathway or mechanism is the trigger of the prolonged dormancy observed. A possible explanation could lie in embryo development before stratification. Embryo morphology has strong effects on dormancy and germination, and seeds with morphological dormancy require a period of ripening or embryo maturation prior to germination (Forbis *et al.* 2002). *Rhinanthus* seeds collected directly fruit ripening have a low germination percentage (Ter Borg 2005), suggesting that these also need a period of after-ripening. Marin et al. (2019) found that *R. minor* seeds from seed lots with smaller embryos (relative to endosperm size) germinated more slowly. If F_1a_ hybrid seeds have smaller embryos that are relatively underdeveloped, this could be a potential mechanism that delays germination. This would require a study of embryo size and gene expression directly after seed formation and during after-ripening.

Studies have revealed that epigenetic process like DNA methylation, histone modifications, and small RNAs play an important role in the different stages of seed development, including embryogenesis, seed maturation, and germination (Greaves *et al.* 2015; Wang and Köhler 2017; Eriksson *et al.* 2020). This contribution of epigenetic regulation has also been reported in the fitness of F_1_ hybrids and heterosis or hybrid vigour (Groszmann *et al.* 2013; Greaves *et al.* 2015). The dormancy of seed can be effected by epigenetic process: it has been shown in *Arabidopsis* that dormancy levels are inherited from the mother through inactivation of the paternal allele of the promoter region of allantoinase (*ALN*) gene by DNA methylation (Iwasaki *et al.* 2019). It seems likely that internal factors, including plant hormones and embryo development, in combination with epigenetic changes may explain the difference in germination behaviour of the *Rhinanthus* reciprocal F_1_ hybrids.

## Supplementary data

Supplementary data are available online at https://academic.oup.com/aob and consist of the following. Figure S1: the length distribution of assembled transcriptomes of F1 seeds. Figure S2: expression percentage by N50 (ExN50) calculated as a fraction of the total expressed data (Ex). Figure S3: GO pathway enrichment analysis based on upregulated genes in F1a_60_NG resulted from comparison of F1a_40_NG vs F1a_60_NG. Figure S4: KEGG pathway enrichment of DE genes in F1a_60_NG vs F1a_60_G, F1a_40_NG vs F1a_60_G and F1m_40_NG vs F1m_40_G comparisons. Table S1: Number of sequenced reads per samples and number of high quality reads have been used for assembly and DE analysis. Table S2: Significant BLASTp matches of obtained ORFs with other plant species. Table S3: GO terms classification of DE transcripts in F1a_60_NG vs F1a_60_G, F1a_40_NG vs F1a_60_G and F1m_40_NG vs F1m_40_G comparisons. Table S4: KEGG pathway enrichment results of DE genes in F1a_60_NG vs F1a_60_G, F1a_40_NG vs F1a_60_G and F1m_40_NG vs F1m_40_G. Table S5: Number of differentially expressed transcripts in all treatment comparisons

## Acknowledgements

This is publication no. BRC375 of the Biodiversity Research Centre of UCLouvain. Computational resources have been provided by the supercomputing facilities of UCLouvain (CISM/UCL) and the Consortium des Équipements de Calcul Intensif en Fédération Wallonie Bruxelles (CÉCI) funded by the Fonds de la Recherche Scientifique de Belgique (F.R.S.-FNRS) under convention 2.5020.11 and by the Walloon Region.

