## Supplemental for "Transcriptome profiling and comparison of *Rhinanthus major* and *Rhinanthus minor* reciprocal F_1_ hybrids during seed stratification and germination"

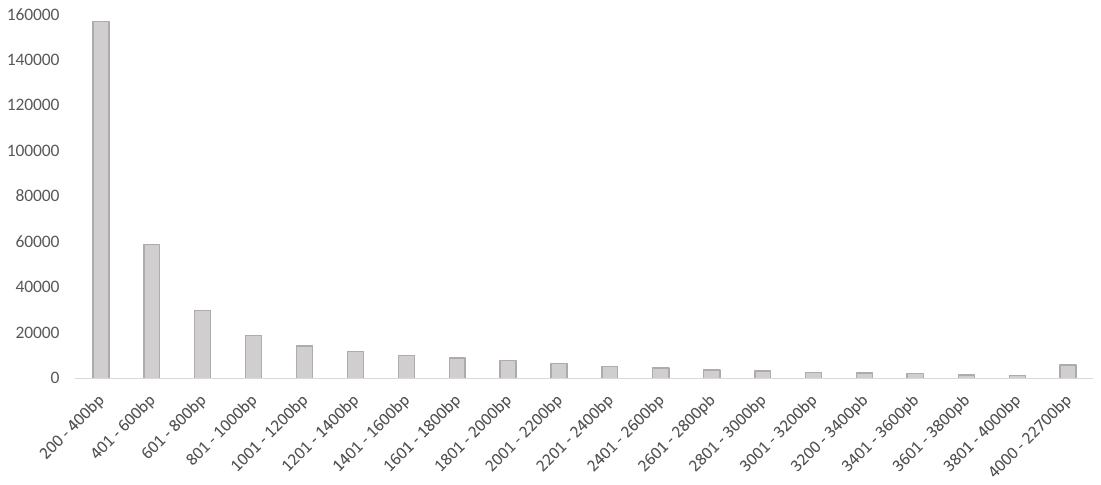

Figure S1. The length distribution of assembled transcriptomes of F_1_ seeds.

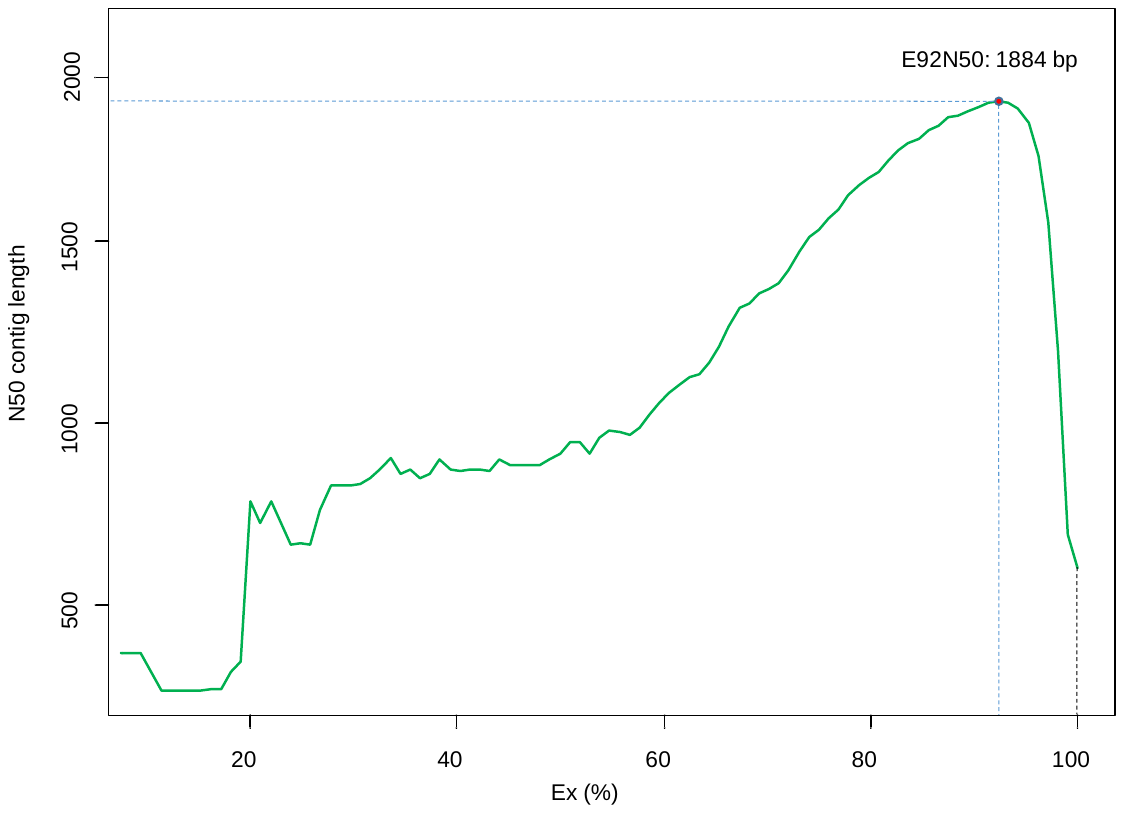

Figure S2. Expression percentage by N50 (ExN50) calculated as a fraction of the total expressed data (Ex). Red dot is ExN50 at the point of assembly saturation (92%) showing that 16,358 transcripts are covered by 92% of reads and N50 of this assembly subset is equal to 1,884 bp.

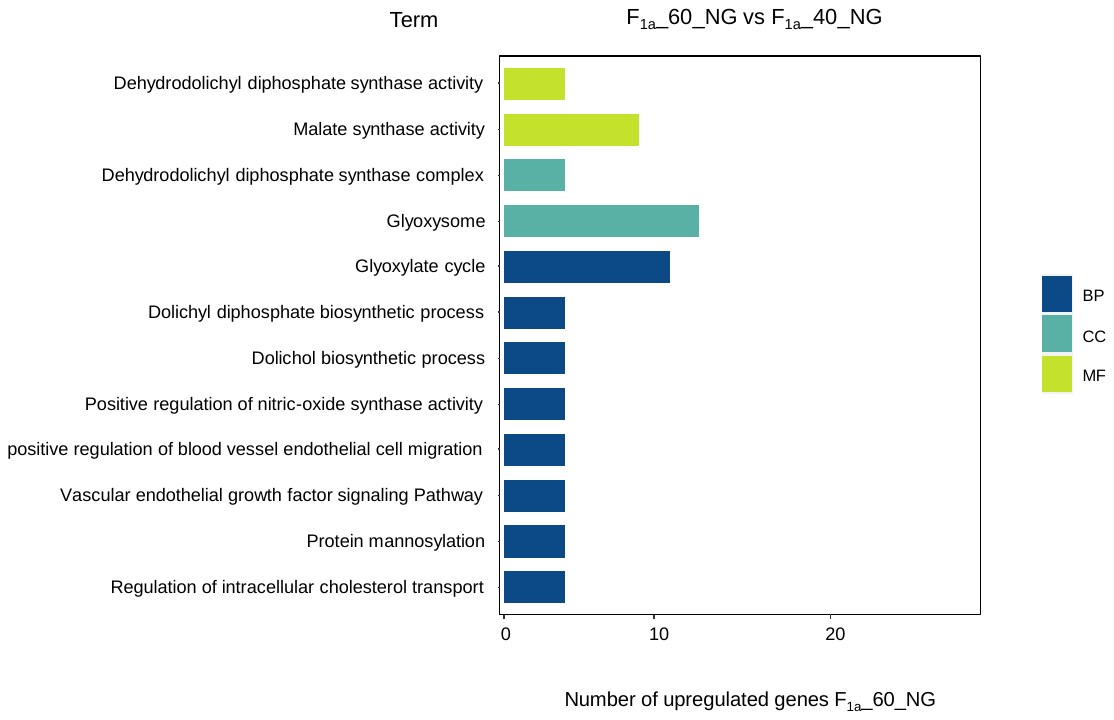

Figure S3. GO pathway enrichment analysis, based on upregulated genes in F_1a__40_NG vs F_1a__60_NG, GO term classification of upregulated genes in F_1a__60_NG. BP = Biological process, CC= Cell cycle, MF= Molecular function

Figure S4A. KEGG pathway enrichment results of DE genes in F1a_60_NG vs F1a_60_G.

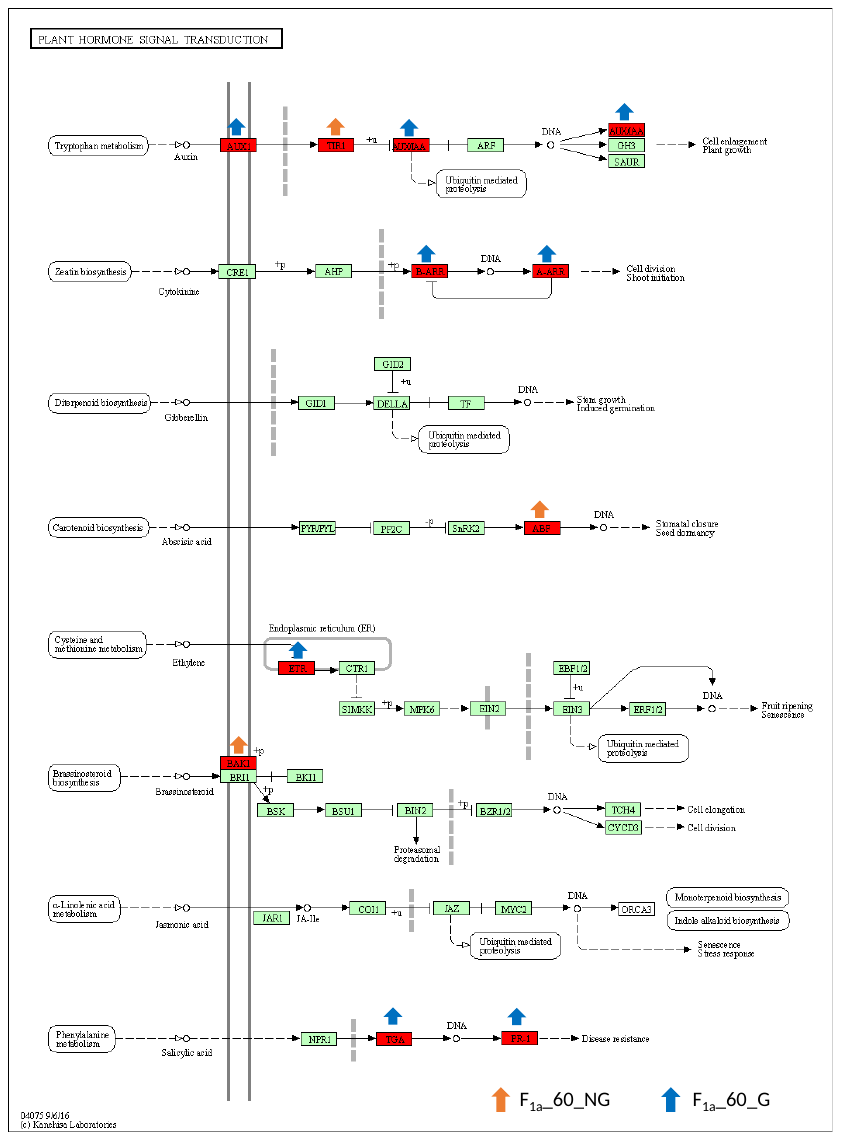

Figure S4B. KEGG pathway enrichment results of DE genes in F1a_40_NG vs F1a_60_G.

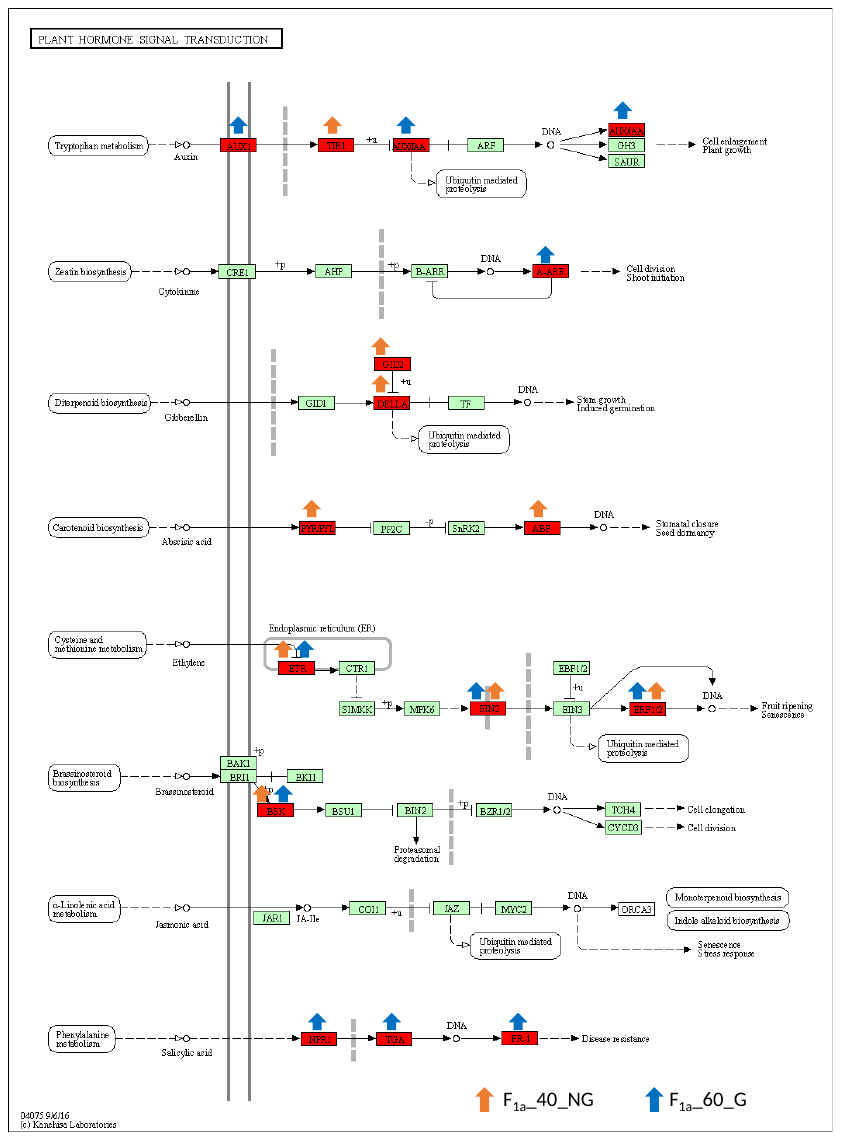

Figure S4C. KEGG pathway enrichment results of DE genes in F1m_40_NG vs F1m_40_G.

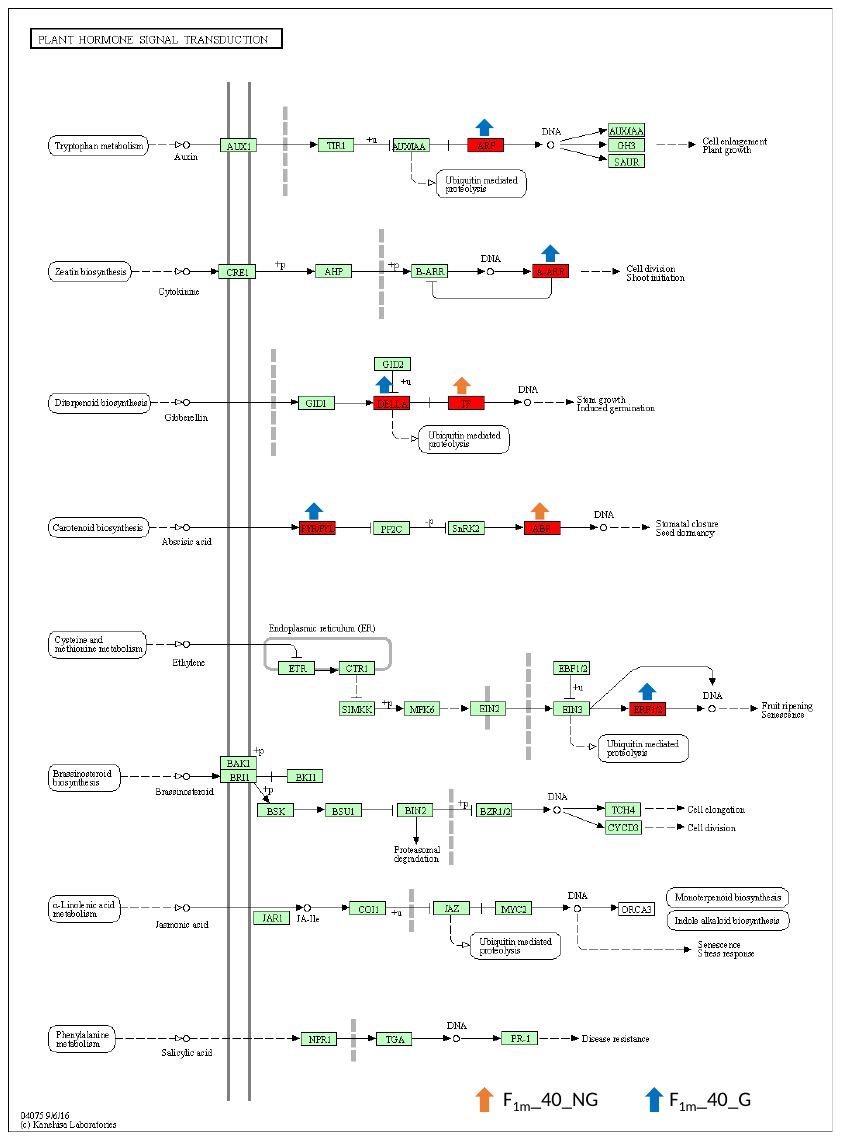

Table S1. Number of sequenced reads per samples details and number of high quality reads (Q>20) have been used for assembly and DE analysis.

| Sample | Raw PE reads | High quality reads | percentage |
| --- | --- | --- | --- |
| F_1a__60_NG_rep1 | 19439586 | 17020996 | 87.55% |
| F_1a__60_NG_rep2 | 21555530 | 18653980 | 86.53% |
| F_1a__60_NG_rep3 | 19523841 | 17641208 | 90.35% |
| F_1a__60_G_rep1 | 18970053 | 16856742 | 88.85% |
| F_1a__60_G_rep2 | 18985001 | 17244304 | 90.83% |
| F_1a__60_G_rep3 | 19472891 | 17531514 | 90.03% |
| F_1a__40_NG_rep1 | 20137400 | 18533775 | 92.03% |
| F_1a__40_NG_rep2 | 19132134 | 17054228 | 89.13% |
| F_1a__40_NG_rep3 | 19434401 | 17137427 | 88.18% |
| F_1m__40_G_rep1 | 22805432 | 20163127 | 88.41% |
| F_1m__40_G_rep2 | 18919276 | 16697648 | 88.25% |
| F_1m__40_G_rep3 | 19025330 | 16603091 | 87.26% |
| F_1m__40_NG_rep1 | 20383643 | 17931419 | 87.96% |
| F_1m__40_NG_rep2 | 19130607 | 16871810 | 88.19% |
| F_1m__40_NG_rep3 | 19297860 | 17064342 | 88.42% |

Table S2. Significant BLASTp matches of *Rhinanthus* hybrid seed ORFs with other plant species in the Plant NR database. Significant with a BitScore>100.

| Species | Family | Match number | Parentage |
| --- | --- | --- | --- |
| *Capsicum annuum* | Solanaceae | 5059 | 5.02% |
| *Gossypium raimondii* | Malvaceae | 4552 | 4.52% |
| *Cynara cardunculus* | Asteraceae | 3737 | 3.71% |
| *Vitis vinifera* | Vitaceae | 3517 | 3.49% |
| *Juglans regia* | Juglandaceae | 3511 | 3.48% |
| *Lupinus angustifolius* | Fabaceae | 3476 | 3.45% |
| *Arabidopsis thaliana* | Brassicaceae | 3308 | 3.28% |
| *Actinidia chinensis* | Actinidiaceae | 2718 | 2.7% |
| *Vitis riparia* | Vitaceae | 2664 | 2.64% |
| *Noccaea caerulescens* | Brassicaceae | 2284 | 2.27% |
| *Citrus clementina* | Rutaceae | 2181 | 2.16% |
| *Striga asiatica* | Orobanchaceae | 2157 | 2.14% |
| Others | - | 61611 | 61.14% |

Table S3A. GO terms classification of DE transcripts in F_1a__60_NG vs F_1a__60_G.

| Category | GO ID | Term | Number of genes | P-Value |
| --- | --- | --- | --- | --- |
| MF | GO:0045735 | nutrient reservoir activity | 43 | 6.15E-22 |
| CC | GO:0005576 | extracellular region | 140 | 2.11E-14 |
| MF | GO:0004497 | monooxygenase activity | 34 | 5.98E-09 |
| MF | GO:0004601 | peroxidase activity | 27 | 7.50E-08 |
| BP | GO:0002253 | activation of immune response | 14 | 2.12E-07 |
| CC | GO:0009505 | plant-type cell wall | 40 | 4.41E-07 |
| BP | GO:0055114 | oxidation-reduction process | 149 | 6.40E-07 |
| BP | GO:0005975 | carbohydrate metabolic process | 82 | 8.62E-07 |
| CC | GO:0048046 | apoplast | 57 | 1.57E-06 |
| MF | GO:0020037 | heme binding | 54 | 1.95E-06 |
| MF | GO:0016705 | oxidoreductase activity, acting on paired donors, with incorporation or reduction of molecular oxygen | 36 | 3.13E-06 |
| CC | GO:0005618 | cell wall | 80 | 1.04E-05 |
| BP | GO:0019253 | reductive pentose-phosphate cycle | 18 | 2.97E-05 |
| BP | GO:0045490 | pectin catabolic process | 17 | 2.97E-05 |
| BP | GO:2000039 | regulation of trichome morphogenesis | 10 | 7.38E-05 |
| MF | GO:0004802 | transketolase activity | 7 | 7.49E-05 |
| BP | GO:0031640 | killing of cells of other organism | 10 | 0.000139 |
| BP | GO:0010731 | protein glutathionylation | 9 | 0.000142 |
| MF | GO:0051920 | peroxiredoxin activity | 10 | 0.000142 |
| MF | GO:0016491 | oxidoreductase activity | 91 | 0.00015 |
| MF | GO:0005506 | iron ion binding | 45 | 0.000161 |
| MF | GO:0004553 | hydrolase activity, hydrolyzing O-glycosyl compounds | 42 | 0.000175 |
| MF | GO:0045330 | aspartyl esterase activity | 13 | 0.000175 |
| MF | GO:0030599 | pectinesterase activity | 14 | 0.000262 |
| BP | GO:0042545 | cell wall modification | 14 | 0.000343 |
| MF | GO:0051119 | sugar transmembrane transporter activity | 10 | 0.000397 |
| BP | GO:0009415 | response to water | 8 | 0.000401 |
| MF | GO:0052716 | hydroquinone:oxygen oxidoreductase activity | 13 | 0.000737 |
| BP | GO:0009938 | negative regulation of gibberellic acid mediated signaling pathway | 7 | 0.000737 |
| BP | GO:0009739 | response to gibberellin | 23 | 0.000941 |
| BP | GO:0046274 | lignin catabolic process | 7 | 0.001028 |
| BP | GO:0010597 | green leaf volatile biosynthetic process | 7 | 0.001341 |
| MF | GO:0004857 | enzyme inhibitor activity | 14 | 0.001521 |
| MF | GO:0070524 | 11-beta-hydroxysteroid dehydrogenase (NADP+) activity | 5 | 0.001585 |
| BP | GO:0006873 | cellular ion homeostasis | 4 | 0.001834 |
| MF | GO:0047216 | inositol 3-alpha-galactosyltransferase activity | 7 | 0.002005 |
| BP | GO:0009664 | plant-type cell wall organization | 18 | 0.002352 |
| MF | GO:0016984 | ribulose-bisphosphate carboxylase activity | 7 | 0.00263 |
| CC | GO:0016021 | integral component of membrane | 429 | 0.00278 |
| MF | GO:0033879 | acetylajmaline esterase activity | 5 | 0.003097 |
| MF | GO:0046910 | pectinesterase inhibitor activity | 10 | 0.00336 |
| BP | GO:0009651 | response to salt stress | 70 | 0.004285 |
| MF | GO:0016711 | flavonoid 3'-monooxygenase activity | 4 | 0.00449 |
| BP | GO:0034220 | ion transmembrane transport | 7 | 0.005876 |
| BP | GO:0009820 | alkaloid metabolic process | 9 | 0.006347 |
| MF | GO:0016985 | mannan endo-1,4-beta-mannosidase activity | 7 | 0.006763 |
| MF | GO:0004322 | ferroxidase activity | 9 | 0.006935 |
| BP | GO:0033619 | membrane protein proteolysis | 10 | 0.012085 |
| MF | GO:0015250 | water channel activity | 11 | 0.014284 |
| BP | GO:0055085 | transmembrane transport | 78 | 0.015211 |
| MF | GO:0019863 | IgE binding | 6 | 0.015291 |
| MF | GO:0016165 | linoleate 13S-lipoxygenase activity | 7 | 0.015291 |
| CC | GO:0009527 | plastid outer membrane | 7 | 0.018428 |
| BP | GO:0051603 | proteolysis involved in cellular protein catabolic process | 22 | 0.020257 |
| MF | GO:0070492 | oligosaccharide binding | 4 | 0.020462 |
| MF | GO:0048030 | disaccharide binding | 4 | 0.020462 |
| MF | GO:0050247 | raucaffricine beta-glucosidase activity | 5 | 0.020534 |
| MF | GO:0050506 | vomilenine glucosyltransferase activity | 5 | 0.020534 |
| BP | GO:0009821 | alkaloid biosynthetic process | 6 | 0.021724 |
| CC | GO:0030875 | rDNA protrusion | 5 | 0.023578 |
| BP | GO:0006952 | defense response | 77 | 0.029063 |
| MF | GO:0015081 | sodium ion transmembrane transporter activity | 17 | 0.031368 |
| BP | GO:0009740 | gibberellic acid mediated signaling pathway | 5 | 0.031368 |
| MF | GO:0045544 | gibberellin 20-oxidase activity | 4 | 0.034726 |
| MF | GO:0102911 | (-)-secoisolariciresinol dehydrogenase activity | 4 | 0.036839 |
| BP | GO:0042744 | hydrogen peroxide catabolic process | 17 | 0.036839 |
| MF | GO:0035673 | oligopeptide transmembrane transporter activity | 7 | 0.039839 |
| BP | GO:0010262 | somatic embryogenesis | 12 | 0.039839 |
| MF | GO:0030742 | GTP-dependent protein binding | 4 | 0.039839 |
| MF | GO:0080115 | myosin XI tail binding | 4 | 0.039839 |
| BP | GO:0048316 | seed development | 21 | 0.044253 |

Table S3B. GO terms classification of DE transcripts in F_1a__40_NG vs F_1a__60_G.

| Category | GO ID | Term | Number of genes | P-Value |
| --- | --- | --- | --- | --- |
| BP | GO:0055114 | oxidation-reduction process | 136 | 5.66E-13 |
| MF | GO:0008422 | beta-glucosidase activity | 37 | 2.21E-08 |
| BP | GO:0071456 | cellular response to hypoxia | 26 | 2.21E-08 |
| MF | GO:0004022 | alcohol dehydrogenase (NAD+) activity | 11 | 8.35E-08 |
| BP | GO:0009061 | anaerobic respiration | 10 | 3.55E-07 |
| MF | GO:0004553 | hydrolase activity, hydrolyzing O-glycosyl compounds | 39 | 7.51E-07 |
| MF | GO:0015250 | water channel activity | 15 | 2.59E-06 |
| BP | GO:0034059 | response to anoxia | 8 | 2.66E-06 |
| CC | GO:0005576 | extracellular region | 90 | 4.84E-06 |
| MF | GO:0016208 | AMP binding | 10 | 2.21E-05 |
| MF | GO:0045330 | aspartyl esterase activity | 12 | 5.41E-05 |
| MF | GO:0030599 | pectinesterase activity | 13 | 5.50E-05 |
| BP | GO:0006952 | defense response | 60 | 5.54E-05 |
| BP | GO:0005975 | carbohydrate metabolic process | 15 | 5.54E-05 |
| BP | GO:0045490 | pectin catabolic process | 71 | 5.54E-05 |
| BP | GO:0042545 | cell wall modification | 41 | 6.14E-05 |
| MF | GO:0020037 | heme binding | 13 | 6.14E-05 |
| MF | GO:0004601 | peroxidase activity | 18 | 7.67E-05 |
| MF | GO:0004737 | pyruvate decarboxylase activity | 6 | 9.31E-05 |
| BP | GO:0002253 | activation of immune response | 11 | 0.00012 |
| MF | GO:0102483 | scopolin beta-glucosidase activity | 17 | 0.000156 |
| MF | GO:0016702 | oxidoreductase activity, acting on single donors with incorporation of molecular oxygen, incorporation of two atoms of oxygen | 12 | 0.000311 |
| MF | GO:0004857 | enzyme inhibitor activity | 13 | 0.000315 |
| BP | GO:0018171 | peptidyl-cysteine oxidation | 7 | 0.000362 |
| MF | GO:0102311 | 8-hydroxygeraniol dehydrogenase activity | 8 | 0.000396 |
| MF | GO:0015267 | channel activity | 12 | 0.000399 |
| BP | GO:0070483 | detection of hypoxia | 7 | 0.000584 |
| MF | GO:0004568 | chitinase activity | 9 | 0.000673 |
| BP | GO:0006032 | chitin catabolic process | 9 | 0.000821 |
| MF | GO:0016705 | oxidoreductase activity, acting on paired donors, with incorporation or reduction of molecular oxygen | 25 | 0.000821 |
| BP | GO:0009415 | response to water | 7 | 0.001391 |
| MF | GO:0016831 | carboxy-lyase activity | 7 | 0.001391 |
| CC | GO:0005618 | cell wall | 9 | 0.001596 |
| MF | GO:0046910 | pectinesterase inhibitor activity | 8 | 0.001596 |
| BP | GO:0016998 | cell wall macromolecule catabolic process | 59 | 0.001596 |
| BP | GO:0051938 | L-glutamate import | 5 | 0.002095 |
| MF | GO:0004190 | aspartic-type endopeptidase activity | 30 | 0.002778 |
| MF | GO:0017172 | cysteine dioxygenase activity | 7 | 0.002834 |
| MF | GO:0008061 | chitin binding | 8 | 0.004017 |
| CC | GO:0099503 | secretory vesicle | 22 | 0.005651 |
| MF | GO:0010333 | terpene synthase activity | 6 | 0.005883 |
| MF | GO:0004497 | monooxygenase activity | 18 | 0.008055 |
| BP | GO:0000272 | polysaccharide catabolic process | 9 | 0.008654 |
| BP | GO:2000039 | regulation of trichome morphogenesis | 6 | 0.009492 |
| MF | GO:0030976 | thiamine pyrophosphate binding | 8 | 0.010214 |
| MF | GO:0038023 | signaling receptor activity | 10 | 0.012523 |
| MF | GO:0033879 | acetylajmaline esterase activity | 4 | 0.01324 |
| BP | GO:0009808 | lignin metabolic process | 4 | 0.01674 |
| BP | GO:0055085 | transmembrane transport | 56 | 0.017794 |
| MF | GO:0004807 | triose-phosphate isomerase activity | 6 | 0.018038 |
| CC | GO:0048046 | apoplast | 37 | 0.018391 |
| MF | GO:0004802 | transketolase activity | 4 | 0.021105 |
| MF | GO:0102406 | omega-hydroxypalmitate O-sinapoyl transferase activity | 3 | 0.021122 |
| MF | GO:0050734 | hydroxycinnamoyltransferase activity | 3 | 0.021122 |
| MF | GO:0016157 | sucrose synthase activity | 6 | 0.02118 |
| BP | GO:0015074 | DNA integration | 20 | 0.025444 |
| CC | GO:0042597 | periplasmic space | 7 | 0.025853 |
| MF | GO:0051213 | dioxygenase activity | 17 | 0.027416 |
| MF | GO:0004519 | endonuclease activity | 21 | 0.031275 |
| BP | GO:0016114 | terpenoid biosynthetic process | 7 | 0.037845 |
| MF | GO:0004619 | phosphoglycerate mutase activity | 3 | 0.039305 |
| MF | GO:0030151 | molybdenum ion binding | 6 | 0.041007 |
| MF | GO:0004351 | glutamate decarboxylase activity | 3 | 0.042743 |
| BP | GO:0010262 | somatic embryogenesis | 60 | 0.044069 |
| MF | GO:0016491 | oxidoreductase activity | 5 | 0.044069 |
| BP | GO:0010345 | suberin biosynthetic process | 6 | 0.044069 |
| BP | GO:0009664 | plant-type cell wall organization | 5 | 0.046025 |
| MF | GO:0015189 | L-lysine transmembrane transporter activity | 13 | 0.046025 |
| BP | GO:0006869 | lipid transport | 18 | 0.046416 |
| BP | GO:0042744 | hydrogen peroxide catabolic process | 14 | 0.049142 |

Table S3C. GO terms classification of DE transcripts in F_1m__40_NG vs F_1m__40_G.

| Category | GO ID | Term | Number of genes | P-Value |
| --- | --- | --- | --- | --- |
| BP | GO:0055114 | oxidation-reduction process | 303 | 3.43E-11 |
| CC | GO:0048046 | apoplast | 121 | 4.67E-11 |
| BP | GO:0002253 | activation of immune response | 239 | 3.91E-10 |
| CC | GO:0005576 | extracellular region | 24 | 3.91E-10 |
| MF | GO:0004553 | hydrolase activity, hydrolyzing O-glycosyl compounds | 89 | 6.93E-10 |
| MF | GO:0004497 | monooxygenase activity | 51 | 1.88E-08 |
| MF | GO:0016491 | oxidoreductase activity | 186 | 1.19E-07 |
| MF | GO:0045735 | nutrient reservoir activity | 38 | 1.97E-07 |
| BP | GO:0009415 | response to water | 15 | 1.98E-07 |
| CC | GO:0005618 | cell wall | 38 | 2.12E-06 |
| MF | GO:0004601 | peroxidase activity | 160 | 2.12E-06 |
| BP | GO:0045490 | pectin catabolic process | 29 | 3.71E-06 |
| BP | GO:0006952 | defense response | 174 | 3.88E-06 |
| MF | GO:0051119 | sugar transmembrane transporter activity | 22 | 5.12E-06 |
| MF | GO:0045330 | aspartyl esterase activity | 17 | 5.12E-06 |
| BP | GO:0031408 | oxylipin biosynthetic process | 30 | 6.35E-06 |
| MF | GO:0051920 | peroxiredoxin activity | 16 | 1.42E-05 |
| BP | GO:0005975 | carbohydrate metabolic process | 141 | 6.34E-05 |
| MF | GO:0020037 | heme binding | 86 | 9.02E-05 |
| MF | GO:0030599 | pectinesterase activity | 22 | 0.000117 |
| MF | GO:0004474 | malate synthase activity | 14 | 0.000168 |
| BP | GO:0042545 | cell wall modification | 22 | 0.000168 |
| BP | GO:0019752 | carboxylic acid metabolic process | 13 | 0.000396 |
| MF | GO:0050614 | delta24-sterol reductase activity | 36 | 0.000494 |
| BP | GO:0010200 | response to chitin | 6 | 0.000494 |
| CC | GO:0009514 | glyoxysome | 41 | 0.00056 |
| MF | GO:0016747 | transferase activity, transferring acyl groups other than amino-acyl groups | 33 | 0.000633 |
| MF | GO:0016984 | ribulose-bisphosphate carboxylase activity | 11 | 0.000633 |
| MF | GO:0004857 | enzyme inhibitor activity | 23 | 0.000683 |
| CC | GO:0009505 | plant-type cell wall | 55 | 0.001497 |
| MF | GO:0016630 | protochlorophyllide reductase activity | 8 | 0.001584 |
| MF | GO:0004740 | pyruvate dehydrogenase (acetyl-transferring) kinase activity | 6 | 0.001807 |
| BP | GO:1904183 | negative regulation of pyruvate dehydrogenase activity | 6 | 0.001807 |
| BP | GO:0055085 | transmembrane transport | 147 | 0.002214 |
| MF | GO:0016762 | xyloglucan:xyloglucosyl transferase activity | 19 | 0.002369 |
| MF | GO:0016705 | oxidoreductase activity, acting on paired donors, with incorporation or reduction of molecular oxygen | 48 | 0.003378 |
| MF | GO:0052716 | hydroquinone:oxygen oxidoreductase activity | 8 | 0.003479 |
| CC | GO:0005615 | extracellular space | 36 | 0.003806 |
| MF | GO:0050661 | NADP binding | 4 | 0.003899 |
| BP | GO:0000717 | nucleotide-excision repair, DNA duplex unwinding | 4 | 0.003899 |
| BP | GO:0000019 | regulation of mitotic recombination | 35 | 0.003899 |
| BP | GO:0009607 | response to biotic stimulus | 17 | 0.004762 |
| BP | GO:0009664 | plant-type cell wall organization | 29 | 0.004851 |
| BP | GO:0046274 | lignin catabolic process | 8 | 0.005074 |
| MF | GO:0046910 | pectinesterase inhibitor activity | 15 | 0.005555 |
| BP | GO:0006097 | glyoxylate cycle | 33 | 0.005921 |
| BP | GO:0009695 | jasmonic acid biosynthetic process | 20 | 0.007218 |
| CC | GO:0016020 | membrane | 312 | 0.007263 |
| MF | GO:0016985 | mannan endo-1,4-beta-mannosidase activity | 6 | 0.00874 |
| MF | GO:0016207 | 4-coumarate-CoA ligase activity | 10 | 0.00874 |
| MF | GO:0016165 | linoleate 13S-lipoxygenase activity | 13 | 0.009916 |
| BP | GO:0009820 | alkaloid metabolic process | 11 | 0.009916 |
| BP | GO:0044262 | cellular carbohydrate metabolic process | 6 | 0.012398 |
| MF | GO:0004351 | glutamate decarboxylase activity | 5 | 0.012991 |
| BP | GO:0048316 | seed development | 39 | 0.013564 |
| MF | GO:0015250 | water channel activity | 18 | 0.013564 |
| BP | GO:0006073 | cellular glucan metabolic process | 13 | 0.014109 |
| BP | GO:0009861 | jasmonic acid and ethylene-dependent systemic resistance | 12 | 0.01424 |
| MF | GO:0033879 | acetylajmaline esterase activity | 6 | 0.014656 |
| MF | GO:0050275 | scopoletin glucosyltransferase activity | 5 | 0.017972 |
| BP | GO:0006979 | response to oxidative stress | 66 | 0.019534 |
| BP | GO:2000899 | xyloglucan catabolic process | 11 | 0.024482 |
| BP | GO:0085030 | symbiotic process benefiting host | 11 | 0.024482 |
| MF | GO:0050506 | vomilenine glucosyltransferase activity | 7 | 0.025211 |
| MF | GO:0050247 | raucaffricine beta-glucosidase activity | 7 | 0.025211 |
| CC | GO:0099503 | secretory vesicle | 45 | 0.026316 |
| BP | GO:0042593 | glucose homeostasis | 6 | 0.028302 |
| BP | GO:0009809 | lignin biosynthetic process | 23 | 0.029063 |
| BP | GO:0045951 | positive regulation of mitotic recombination | 4 | 0.029063 |
| BP | GO:0071446 | cellular response to salicylic acid stimulus | 6 | 0.029063 |
| BP | GO:0019253 | reductive pentose-phosphate cycle | 20 | 0.029186 |
| MF | GO:0022857 | transmembrane transporter activity | 91 | 0.030283 |
| MF | GO:0005199 | structural constituent of cell wall | 9 | 0.030283 |
| MF | GO:0030598 | rRNA N-glycosylase activity | 6 | 0.030581 |
| MF | GO:0008171 | O-methyltransferase activity | 11 | 0.032585 |
| CC | GO:0042807 | central vacuole | 6 | 0.033455 |
| MF | GO:0016758 | transferase activity, transferring hexosyl groups | 33 | 0.038483 |
| MF | GO:0070006 | metalloaminopeptidase activity | 10 | 0.038666 |
| BP | GO:0015824 | proline transport | 4 | 0.038999 |
| BP | GO:0006829 | zinc ion transport | 7 | 0.043737 |
| BP | GO:0009741 | response to brassinosteroid | 19 | 0.049457 |

Table S4. KEGG pathway enrichment results of DE genes in F_1a__60_NG vs F_1a__60_G, F_1a__40_NG vs F_1a__60_G and F_1m__40_NG vs F_1m__40_G.

| Term | ID | Number of genes | P-Value |
| --- | --- | --- | --- |
| **F_1a__60_NG vs F_1a__60_G** |  |  |  |
| Metabolic pathways | ath01100 | 47 | 0.000298 |
| Ubiquitin mediated proteolysis | ath04120 | 8 | 0.004205 |
| Plant hormone signal transduction | ath04075 | 11 | 0.004205 |
| Biosynthesis of secondary metabolites | ath01110 | 25 | 0.005014 |
| Amino sugar and nucleotide sugar metabolism | ath00520 | 6 | 0.04045 |
| **F_1a__40_NG vs F_1a__60_G** |  |  |  |
| Metabolic pathways | ath01100 | 105 | 1.33E-12 |
| Plant hormone signal transduction | ath04075 | 23 | 2.26E-06 |
| Glycerolipid metabolism | ath00561 | 10 | 4.5E-05 |
| Biosynthesis of secondary metabolites | ath01110 | 46 | 0.00027 |
| Glycerophospholipid metabolism | ath00564 | 10 | 0.001224 |
| Amino sugar and nucleotide sugar metabolism | ath00520 | 11 | 0.002452 |
| Plant-pathogen interaction | ath04626 | 12 | 0.004828 |
| alpha-Linolenic acid metabolism | ath00592 | 6 | 0.005911 |
| MAPK signaling pathway - plant | ath04016 | 10 | 0.007646 |
| Ubiquitin mediated proteolysis | ath04120 | 10 | 0.010951 |
| Glycosphingolipid biosynthesis - globo and isoglobo series | ath00603 | 3 | 0.013633 |
| Glutathione metabolism | ath00480 | 8 | 0.014248 |
| Fatty acid biosynthesis | ath00061 | 5 | 0.020304 |
| Fatty acid degradation | ath00071 | 5 | 0.026145 |
| Fatty acid metabolism | ath01212 | 6 | 0.026145 |
| Fatty acid elongation | ath00062 | 4 | 0.044855 |
| Circadian rhythm - plant | ath04712 | 4 | 0.044855 |
| Arginine biosynthesis | ath00220 | 4 | 0.044855 |
| **F_1m__40_NG vs F_1m__40_G** |  |  |  |
| Metabolic pathways | ath01100 | 43 | 6.93E-08 |
| Biosynthesis of secondary metabolites | ath01110 | 19 | 0.00694 |
| alpha-Linolenic acid metabolism | ath00592 | 4 | 0.00694 |
| Plant hormone signal transduction | ath04075 | 8 | 0.011712 |
| Circadian rhythm - plant | ath04712 | 3 | 0.024828 |
| Linoleic acid metabolism | ath00591 | 2 | 0.024828 |
| MAPK signaling pathway - plant | ath04016 | 5 | 0.024828 |
| Glycerophospholipid metabolism | ath00564 | 4 | 0.041279 |
| Starch and sucrose metabolism | ath00500 | 5 | 0.045252 |

Table S5. Number of differentially expressed transcripts in all treatment comparisons ([FDR] < 0.05 and |log2FC| ≥ 2) in F_1_ hybrids between *Rhinanthus major* (a) and *R. minor* (m), the letter indicating the maternal parent. NG = non-germinated seeds, G = germinated seeds, after 40 or 60 days on wet filter paper at 4°C.

|  | F_1a__60_NG | F_1a__60_G | F_1a__40_NG | F_1m__40_NG |
| --- | --- | --- | --- | --- |
| F_1a__60_NG | - |  |  |  |
| F_1a__60_G | 1893 | - |  |  |
| F_1a__40_NG | 2416 | 4165 | - |  |
| F_1m__40_NG | 4421 | 5189 | 6052 | - |
| F_1m__40_G | 8096 | 5762 | 8969 | 1420 |
